# Relaxed constraint and thermal desensitization of the cold-sensing ion channel TRPM8 in mammoths

**DOI:** 10.1101/397356

**Authors:** Sravanthi Chigurapati, Mike Sulak, Webb Miller, Vincent J. Lynch

## Abstract

Unlike the living elephants, which live in warm tropical and subtropical habitats, mammoths lived in extreme cold environments where average winter temperatures ranged from −30° to −50°C. Like other cold adapted and artic species, mammoths evolved of suite of morphological and molecular adaptations that facilitated life in the cold. Here we reanalyze mammoth genomes and find that genes with mammoth-specific amino acid substitutions are enriched in functions related to temperature sensation. Among the genes with mammoth-specific amino acid substitutions is TRPM8, which mediates sensitivity to nonnoxious cool temperatures from 25– 28°C and cooling sensations induced by the chemical agonists menthol and icilin. We find that TRPM8 evolved rapidly in the mammoth stem-lineage, likely because of an episode of relaxed purifying selection. Functional characterization of resurrected mammoth and ancestral TRPM8 indicates that the mammoth TRPM8 is desensitized to cold but maintains sensitivity to menthol and icilin. These data suggest that as mammoths evolved into a cold tolerant species they lost the need for a cold-sensitive TRPM8.

## Introduction

Living and extinct elephants originated Africa and dispersed into warm tropical and subtropical habitats in Africa and Eurasia. Woolly mammoths (*Mammuthus primigenius*), in contrast, lived in the extreme cold of the mammoth-steppe where average winter temperatures ranged from −30° to −50°C (MacDonald et al., 2012). Cold adapted and artic species have evolved molecular and physiological adaptations to their circadian systems (Bloch et al., 2013; Lu et al.), adipose biology (Fumagalli et al., 2015; Nelson et al., 2014; Welch et al., 2014), and temperature sensation (Matos-Cruz et al., 2017) to deal with serve cold and long periods of persistent dark in winter and light in summer, suggesting mammoths may have also evolved similar adaptations. While temperature sensation in mammals is a complex sensory process in which temperature-sensitive primary sensory neurons convey thermal information from the skin and peripheral organs to the CNS (Vriens et al., 2014), temperature-sensitive transient receptor potential (thermoTRP) cation channels are the major molecular thermosensors (MacDonald et al., 2012; Vriens et al., 2014).

Temperature-sensitive TRP proteins function as tetrameric channels, which are activated (opened) at specific temperatures to allow calcium influx. The primary thermoTRP that mediates sensitivity to cold is TRPM8, which is generally activated at temperatures less than 26°C *in vivo* and *in vitro*. TRPM8 knockout mice, for example, have profoundly diminished responses to cold and severe behavioral deficits in their ability to discriminate between cold and warm surfaces. Similarly, thirteen-lined ground squirrels (*Ictidomys tridecemlineatus*) and Golden hamsters (*Mesocricetus auratus*) have TRPM8 channels with dramatically reduced cold sensitivity and do not avoid cold temperatures (Bloch et al., 2013; Lu et al.; Matos-Cruz et al., 2017). TRPM8 is also activated in response to cold temperatures and ligands that induce cooling sensations such as menthol, eucalyptol, and the ‘super-cooling’ agent icilin in heterologous expression systems (Fumagalli et al., 2015; Nelson et al., 2014; Vriens et al., 2014; Welch et al., 2014). Remarkably administration TRPM8 antagonists to mice and rats decreased their core body temperature, implicating TRPM8 in maintenance of body temperature in addition to cold sensation (Almeida et al., 2012; Gavva et al., 2012; Matos-Cruz et al., 2017; Reimúndez et al., 2018).

Here we show that genes with mammoth specific amino acid changes are enriched in temperature sensing functions. To infer if mammoths may have evolved adaptations to temperature sensation we reanalyzed African elephant, Asian elephant, and woolly mammoth genomes to identify amino acid substitutions in genes related to temperature sensation, including TRPM8. We identified four amino acid substitutions that occurred in the mammoth stem-lineage, two of which are predicted by PolyPhen-2 to be deleterious. We show that TRPM8 evolved rapidly in the mammoth stem-lineage, but that the signal of rapid evolution is more consistent with relaxed purifying selection rather than positive selection. Finally we functionally characterize the ancestral mammoth TRPM8 protein and show that it is not activated in response to cold but maintains sensitivity to the chemical ligands menthol and icilin. These data suggest that mammoths may have sensed cold temperatures differently than extant elephants and other mammals.

## Results and discussion

### Genes with mammoth-specific amino acid substitutions are enriched in temperature sensing functions

We previously compared two woolly mammoth genomes to three Asian elephant genomes in order to identify genes with mammoth specific amino acid changes (Lynch et al., 2015; Vriens et al., 2014); These genes were enriched in numerous mouse KO phenotypes, most intriguingly ‘abnormal thermal nociception’ (13 genes). In an attempt to replicate and extend these results, we reanalyzed elephant genomes including an African elephant, three Asian elephants, and the Oimyakon (11-fold coverage), M4 (20-fold coverage), and Wrangel Island (17-fold coverage) mammoth genomes; neither the Oimyakon nor Wrangel Island mammoth genomes were included in our original analyses (Lynch et al., 2015). We then compared mammoth genomes to African and Asian elephant genomes and identified 6056 mammoth-specific amino acid changes in 4009 protein coding genes.

As in our previous analyses, we again found that genes with mammoth specific amino acid changes were enriched in numerous mouse knockout (KO) phenotypes including 33 nervous system phenotypes related to temperature sensation (**Fig. 1**) such as ‘abnormal thermal nociception’ (P=1.31×10^-4^, FDR=2.08×10^-^3), ‘abnormal touch/nociception’ (P=2.93×10^-4^, FDR=4.03×10^-3^), and ‘increased thermal nociceptive threshold’ (P=1.58×10^-3^, FDR=1.44×10^-2^).Among the 31 genes with mammoth specific amino acid changes annotated as having ‘abnormal thermal nociception’ phenotypes in KO mice are thermoTRP channels that mediate temperature sensation, including TRPM8 the primary molecular cold sensor (four amino acid changes), TRPA1 a sensor or noxious cold (one amino acid change), and TRPV3 a sensor a warm temperatures (three amino acid changes), as well as the voltage-gated channel SCN10A (Na_v_1.8) which plays a crucial role in the sensation of painful cold (four amino acid changes).

**Figure 1.**
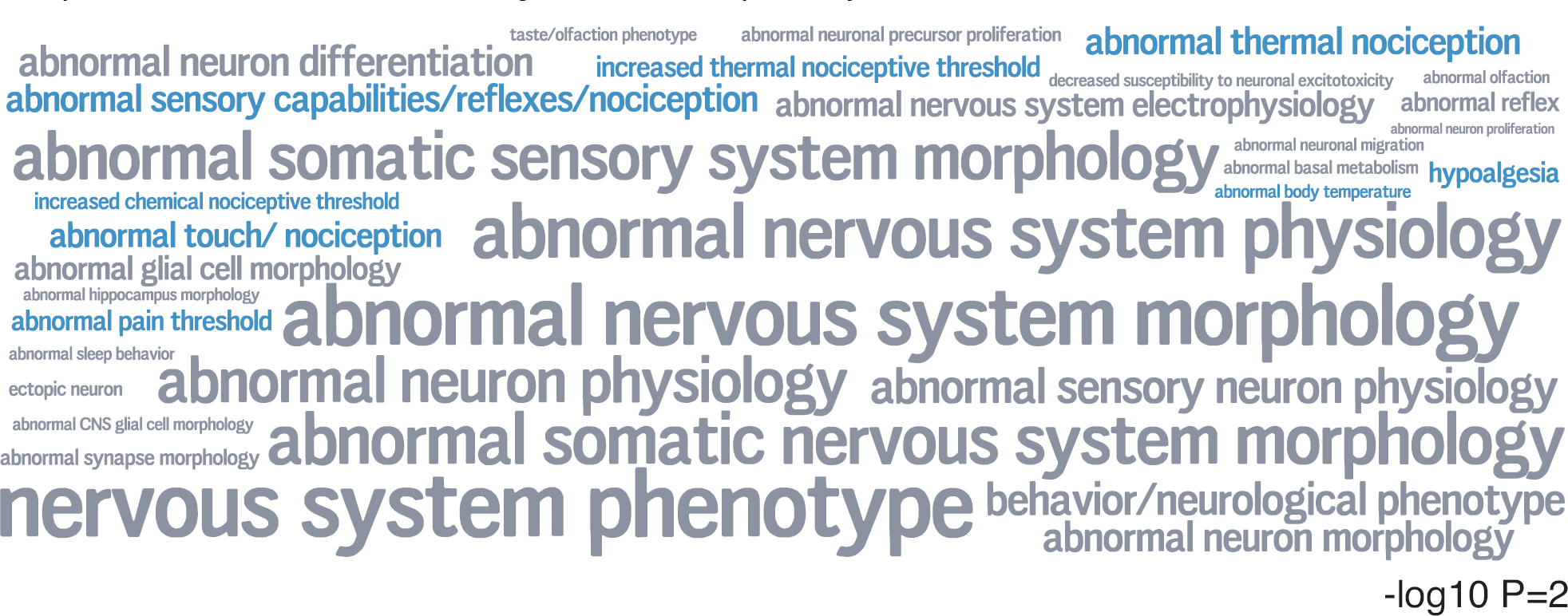
Genes with mammoth-specific amino acid changes are enriched in mouse knockout (KO) phenotypes related to the nervous system and temperature sensation. Word cloud of 33 mouse KO phenotypes related to the nervous system (MP:0003631) enriched among protein-coding genes with mammoth-specific amino acid changes. Phenotype terms are scaled to the −log10 *P*-value of phenotype enrichment (hypergeometric test), inset scale word “−log10=2”. Phenotype terms are color-coded by general nervous system phenotypes, and phenotype related to temperature sensation.

### Rapid evolution of TRPM8 in mammoths

We identified *TRPM8* orthologs from 93 vertebrates, including African savannah elephant (*Loxodonta africana*), African forest elephant (*Loxodonta cyclotis*), straight-tusked elephant (*Palaeoloxodon antiquus*), three Asian elephants (*Elephas maximus*), four Woolly mammoths (*Mammuthus primigenius*), a Columbian mammoth (*Mammuthus columbi*) and an American mastodon (*Mammut americanum*) using previously published genomes (Lynch et al., 2015; Palkopoulou et al., 2018), and used several complementary methods to characterize the strength and direction of selection acting on *TRPM8* in mammoths. We found that the adaptive branch-site random effects likelihood (aBSREL) method (Kosakovsky Pond et al., 2011; Smith et al., 2015), which allows for variable *d*_*N*_*/d*_*S*_ (ω) rates across lineages and sites, inferred a site class with ω>1 in the mammoth stem-lineage but the results were not statistically significant (LRT=2.34, *P*=0.11). We next used a random effects branch-site model of relaxed selection intensity (RELAX) (Wertheim et al., 2015) to test if the episode of rapid evolution in the mammoth stem-lineage was consistent with a signal of relaxed selection intensity (K<1). Consistent with an episode of reduced purifying selection, RELAX inferred significant evidence for a relaxation in the selection intensity in the mammoth stem-lineage (K=0.00; LRT=11.24, *P*=0.001). We also used aBSREL and RELAX to test if the relaxation in the intensity of selection persisted into descendent mammoth lineages, and found that aBSREL inferred single *d*_*N*_*/d*_*S*_rate (ω=0.31) and no evidence relaxed in the selection intensity within the mammoth clade (K=0.12; LRT=3.23, *P*=0.072). These data suggest that the episode of rapid evolution in the *TRPM8* gene in the mammoth stem-lineage resulted from relaxed purifying selection rather than positive selection, and that purifying selection continued in descendent mammoth lineages.

### Mammoth-specific amino acid substitutions in TRPM8 are predicted to be deleterious

We reconstructed the sequence of the Asian elephant/mammoth ancestral TRPM8 (AncGajah) and ancestral mammoth (AncMammoth) TRPM8 proteins, and identified four amino acid substitutions that are fixed within mammoths at sites that are invariant or nearly invariant in 88 other vertebrates (**Fig. 2A**). To infer if these four amino acid substitutions may have had structural or functional consequences, we computationally predicted their functional effects using PolyPhen-2. PolyPhen-2 classified the R368H and C711S substitutions as ‘probably damaging’ whereas the A533S and G710S substitutions were predicted to be ‘benign’. We also inferred structural models of the AncGajah and AncMammoth TRPM8 proteins and found that the G710S and C711S substitutions were clustered in the pre-S1 domain nearby a site previously shown to affect channel function, while R368H and A533S were located in unstructured loops (**Fig. 2B/C**). We used the mCSM method (Pires et al., 2014b) implemented in DUET (Pires et al., 2014a) to predict if these substitutions might effect protein stability and found that the R368H (ΔΔG=-0.679), A533S (ΔΔG=-1.431),G710S(ΔΔG=-0.689),C711S(ΔΔG=-0.283) were all predicted to be destabilizing (sum ΔΔG=-3.761). We also observed that the G710S and C711S substitutions are located in the pre-S1 domain nearby a residue that mediates the sensitivity of TRPM8 to cold. These data suggest that at least some of the mammoth specific substitutions in TRPM8 may have had functional consequences.

**Figure 2.**
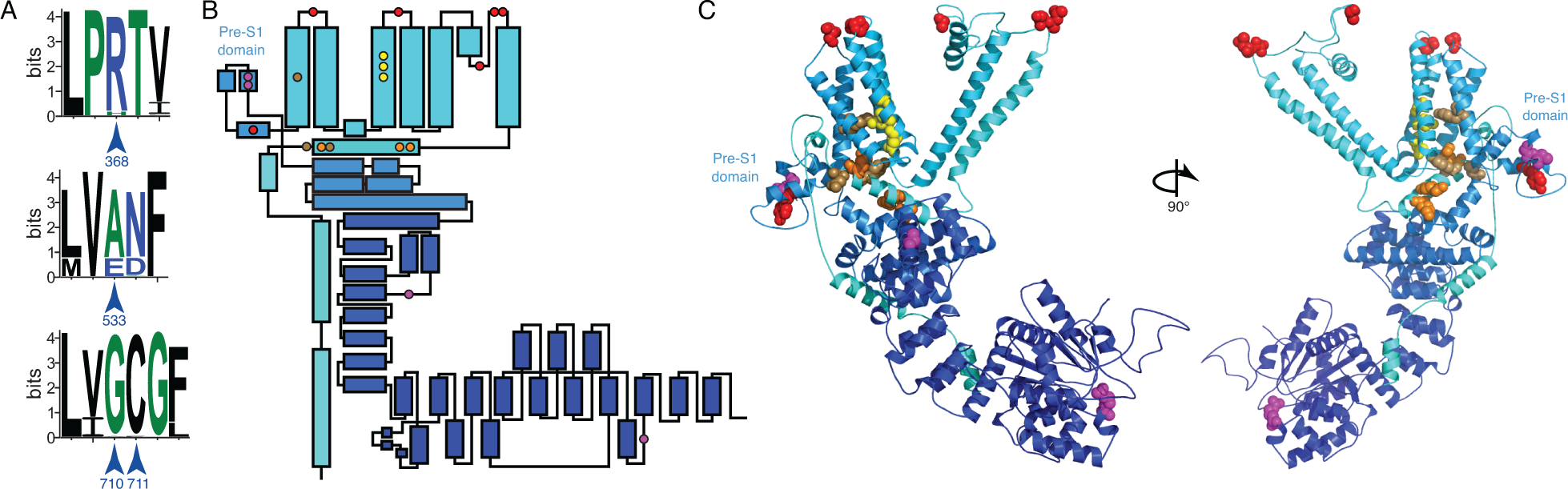
Mammoth-specific amino acid changes in TRPM8 occur near sites that mediate cold responses. **(A)** Sequence logo showing conservation of sites 368 (top), 533 (middle), and 710/711 (bottom) from 88 vertebrate species. **(B)** Topology diagram of TRPM8 showing the location of secondary structural elements, mammoth specific amino acid substitutions, and functional sites. Helices are shown as boxes, mammoth-specific substitutions as magenta circles, resides that mediate responses to cold as red circles, responses to menthol as tan circles, icilin binding as yellow circles, and PIP2 binding as orange circles. **(C)** Homology model of the AncMammoth TRPM8 protein. Coloring of secondary structural elements and functional residues follows panel B. The pre-S1 domain is labeled.

### Thermal desensitization of TRPM8 in mammoths

To functionally characterize the mammoth-specific amino acid substitutions in TRPM8, we resurrected the AncMammoth and AncGajah *TRPM8* genes (**Fig. 3A**) and measured their temperature-dependent activity in transiently transfected HEK293 cells using a Fluo-4 calcium influx assay (Aneiros and Dabrowski, 2009; Reubish et al., 2009). We found that the AncGajah TRPM8 had a similar activation profile as other mammalian TRPM8 proteins, for example, it was activated at ∼22°C, had half maximum activities (T_50_) at ∼17°C, and maximal activities (T_max_) at 2°C (**Fig. 3B**). The AncMammoth TRPM8 protein, however, was almost completely insensitive to cooling and was >90% less active than the AncGajah channel at T_max_ (**Fig. 3B**). Both channels, however, were robustly activated in response to the chemical TRPM8 agonists menthol and icilin (**Fig. 3C**). These data indicate that mammoth-specific amino acid changes specifically affect the ability of TRPM8 to respond to temperature changes but do not alter its sensitivity to chemical agonists.

**Figure 3.**
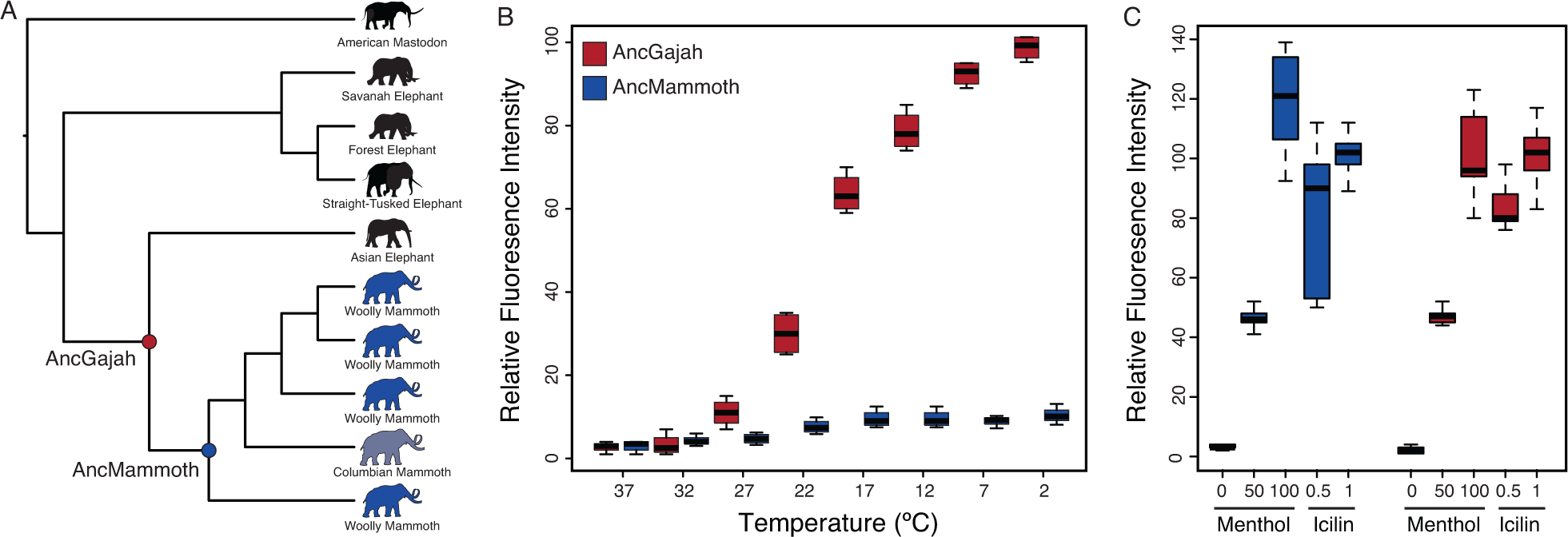
Mammoth TRPM8 is insensitive to cold, but is activated by chemical agonists. **(A)** Phylogenetic relationships between the Proboscideans used for TRPM8 ancestral sequence reconstruction. The AncGajah and AncMammoth nodes are labeled. **(B)** Temperature-dependent activity of AncGajah (red) and AncMammoth (blue) TRPM8 in transiently transfected HEK293 cells assayed using a Fluo-4 calcium influx assay. Data are shown as relative fluorescence intensity standardized to the maximal observed intensity (AncGajah at 2°C). n=36. **(C)** Activity of AncGajah (red) and AncMammoth (blue) TRPM8 in transiently transfected HEK293 cells treated with either menthol (uM) or icilin (uM). Data are shown as relative fluorescence intensity standardized to the maximal observed intensity (AncGajah). Data are shown as relative fluorescence intensity. Menthol and icilin responses are standardized to AncGajah treated with 100uM menthol and 1uM icilin, respectively. n=18.

## Conclusions

Mammoths evolved a suite of adaptations to extreme cold, including molecular adaptations such as cold adapted hemoglobin (Campbell et al., 2010; Yuan et al., 2013; 2011) and reduced sensitivity of the thermoTRP TRPV3 to warm temperatures (Lynch et al., 2015). Here we show that the thermoTRP TRPM8 experienced an episode of relaxed purifying selection in the mammoth stem-lineage and was not activated by cold. These data suggest that selection to maintain a functional cold-activated TRPM8 was lost as mammoths colonized extremely cold environments, and thus lost the need to sense or avoid cold temperatures. While we cannot determine how loss of TRPM8 temperature sensitivity may have effected mammoths, we can infer likely consequences based on mouse TRPM8 knockout (KO) phenotypes as well as the behavior of ground squirrels and Golden hamsters, which have TRPM8 proteins that are insensitive to cold (Matos-Cruz et al., 2017). TRPM8 KO mice, for example, have reduced avoidance of cold temperatures (Bautista et al., 2007; Colburn et al., 2007; Dhaka et al., 2007) as do ground squirrels and Golden hamsters (Matos-Cruz et al., 2017). Remarkably, TRPM8 KO mice also have a robust preference for 5°C over the mildly noxious hot 45°C in thermal tolerance tests suggesting mammoths may have preferred cold to hot temperatures.

## Methods

### Functional Inference of Mammoth-Specific Amino Acid Substitutions

Our previous analyses of woolly mammoth genomes included the M4 and M25 mammoths (Lynch et al., 2015), however, M25 is likely contaminated with multiple mammoth individuals (Rogers and Slatkin, 2017). To replicate and extend these results, therefore, we used the bioinformatics pipeline we previously described (Lynch et al., 2015) to reanalyze genomes of the African elephant, three Asian elephants, and the M4 (Lynch et al., 2015), Oimyakon (11-fold coverage) and Wrangel Island (17-fold coverage) mammoth genomes (Palkopoulou et al., 2015). We thus identified 6056 mammoth-specific amino acid changes in 4009 protein coding genes. Next, we used Vlad (http://proto.informatics.jax.org/prototypes/vlad/) to mine the mouse KO phenotype data at Mouse Genome Informatics (http://www.informatics.jax.org) for the genes with mammoth-specific substitutions and to identify mouse KO phenotypes in which those genes were enriched. The results can be found in Supplementary Tables.

### TRPM8 Ancestral Sequence Reconstruction and Gene Synthesis

To reconstruct the ancestral mammoth (AncMammoth) and Asian elephant/mammoth (AncGajah) TRPM8 protein sequences, we: (1) Identified TRPM8 genes from 93 vertebrate (http://hgdownload.cse.ucsc.edu/goldenPath/hg19/multiz100way), including the African Savannah elephant (*Loxodonta africana*), African Forest elephant (*Loxodonta cyclotis*), Asian elephant (*Elephas maximus*), four woolly mammoths (*Mammuthus primigenius*), Columbian mammoth (*Mammuthus columbi*), and straight-tusked elephant (*Palaeoloxodon antiquus*), and the American Mastodon (*Mammut americanum*) (Palkopoulou et al., 2015; 2018); (2) aligned the translated sequences with MUSCLE (Edgar, 2004); (3) inferred the JTT model as the best-fitting model of amino acid substitution using the model selection module implemented in Datamonkey (Delport et al., 2010); and (4) used joint (Pupko et al., 2000), marginal (Yang et al., 1995), and sampled (Nielsen, 2002) maximum likelihood ancestral sequence reconstruction (ASR) methods implemented in the ASR module of Datamonkey. The JTT model incorporated a general discrete model of site-to-site rate variation with three rate classes. We used the phylogeny associated with the multiz100way alignment, except for the phylogenetic relationships within Proboscideans which was based of the Palkopoulou et al. 2018 phylogeny. We inferred the phylogenetic relationships among mammoths using the TRPM8 alignment and PhyML (Guindon et al., 2010) using the same substitution model described above. Support for the ancestral mammoth and Gajah reconstructions were 1.0 for all sites under joint, marginal, and sampled likelihoods. We also used the Datamonkey sever to test for relaxed constraint and positive selection using aBSREL (Smith et al., 2015) and RELAX (Wertheim et al., 2015).

The AncMammoth and AncGajah TRPM8 genes, including a C-terminal FLAG tag (MDYKDDDDK) and a Kozack sequence incorporated into the initiating methionine (gccaccATGG) were synthesized by GeneScript using human codon usage and cloned into the mammalian expression vector pcDNA5/FRT (Invitrogen).

### TRPM8 Structure Modeling

The AncMammoth and AncGajah protein structures were modeled on the high-resolution cryo-EM structure of TRPM8 (Yin et al., 2018). Initial structural models of the AncMammoth and AncGajah proteins in the were generated using I-TASSER (Roy et al., 2010; Zhang, 2008), structural models were refined with ModRefiner (Xu and Zhang, 2011), using the TRPM8 channel (PDB: 6BPQ) as a reference structure.

### TRPM8 Function Assays

We used the Fluo-4 NW Calcium Assay Kit (Life Technologies) to determine the temperature response of the AncMammoth and AncGajah TRPM8 proteins. HEK293 cells (ATCC CRL-1573), were cultured in MEM supplemented with 10% (v/v) fetal bovine serum (FBS) in a 37°C humidity-controlled incubator with 10% CO2. HEK293 cells growing in 10-cm plates were transiently transfected at 80% confuency with 24 μg expression vector for the AncMammoth or AncGajah TRPM8 genes or empty pcDNA5/FRT using Lipofectamine LTX+ (Life Technologies), using the standard protocol. 48 hr after transfection, cells were harvested by trypsinization, centrifuged, resuspended in Hank’s balanced salt solution (HBSS) and HEPES assay buffer containing 2.5 mM probenecid, and transferred to a 96-well plate at 150,000 cells/well in 50 μl assay buffer. Temperature-dependent calcium influx was assayed using the Fluo-4 NW Calcium Assay Kit (Molecular Probes) and a high-throughout qPCR-based assay (Aneiros and Dabrowski, 2009, Reubish et al., 2009). After an initial 30-min loading at 37°C, the temperature was lowered from 37°C to 2°C in 5°C steps, and fluorescence was measured after 30 s at each temperature using a Bio-Rad CFX-96 real-time PCR machine. Fluo-4 fluorescence was measured using channel 1 (Sybr/FAM). Fluo-4 fluorescence of cells transfected with the AncMammoth or AncGajah TRPM8 genes was normalized by the Fluo-4 fluorescence of empty pcDNA5/FRT-transfected controls. Similarly, we assayed menthol (Sigma) and icilin (Sigma) sensitivity by quantifying Fluo-4 fluorescence of cells transfected with the AncMammoth or AncGajah TRPM8 genes normalized by the Fluo-4 fluorescence of empty pcDNA5/FRT-transfected controls. All temperature experiments included 32 biological replicates and were repeated in three independent experiments. Menthol and icilin experiments included 18 replicates per treatment.

